# The Planarian Anatomy Ontology: A resource to connect data within and across experimental platforms

**DOI:** 10.1101/2020.08.14.251579

**Authors:** Stephanie H. Nowotarski, Erin L. Davies, Sofia M. C. Robb, Eric J. Ross, Nicolas Matentzoglu, Viraj Doddihal, Mol Mir, Melainia McClain, Alejandro Sánchez Alvarado

**Author notes:** denotes equal contribution to work (Stephanie H. Nowotarski, Erin L. Davies, Sofia M. C. Robb).

## Abstract

As the planarian *Schmidtea mediterranea* (*Smed*) gains popularity as a research organism, the need for standard anatomical nomenclature is increasingly apparent. A controlled vocabulary streamlines data annotation, improves data organization, and enhances cross-platform and cross-species searchability. We created the Planarian Anatomy Ontology (PLANA), an extendable framework of defined *Smed* anatomical terms organized using relationships. The most current version contains over 800 terms that describe *Smed* anatomy from subcellular to system-level across all life cycle stages, in intact animals, and regenerating body fragments. Terms from other anatomy ontologies were imported into PLANA to promote ontology interoperability and comparative anatomy studies. To demonstrate the utility of PLANA for data curation, we created web-based resources for planarian embryogenesis, including a staging series and molecular fate mapping atlas, as well as a searchable Planarian Anatomy Gene Expression database, which integrates a variety of published gene expression data and allows retrieval of information of all published sequences associated with specific planarian anatomical regions. Finally, we report methods for continued curation of PLANA, providing a path for expansion and evolution of this community resource.

**Summary Statement:** We report construction of an anatomy ontology for an emerging research organism and show its use to curate and mine data across multiple experimental platforms.

## Introduction

Researchers using the free-living, freshwater planarian *Schmidtea mediterranea (Smed)* are rapidly generating genomic, transcriptomic, phenotypic, and anatomical data. However, the field has neither a standardized vocabulary nor a universally accepted set of gene/transcript models to allow for quick, reliable navigation and integration of data across experimental platforms and publications. *Smed* anatomical information has been garnered using techniques for structural and ultrastructural visualization (e.g., histological staining, scanning and transmission electron microscopy), as well as molecular techniques that report gene expression or protein localization *in situ* (e.g., whole-mount *in situ* hybridization, immunohistochemistry, and immunofluorescence). Gene discovery has been facilitated through sequenced *Smed* genome assemblies (Robb, Ross and Sánchez Alvarado, 2008; Robb *et al*., 2015; Grohme *et al*., 2018), and *de novo* assembled *Smed* transcriptomes (Adamidi *et al*., 2011; Sandmann *et al*., 2011; Labbé *et al*., 2012; Rouhana *et al*., 2012; Srivastava *et al*., 2014; Tu *et al*., 2015; Brandl *et al*., 2016). Microarray analysis (Eisenhoffer, Kang and Sánchez Alvarado, 2008; Wagner, Ho and Reddien, 2012), bulk (Blythe *et al*., 2010; Solana *et al*., 2012; Davies *et al*., 2017) and single-cell RNA-Seq (Wurtzel *et al*., 2015; Fincher *et al*., 2018; Plass *et al*., 2018; Zeng *et al*., 2018) identified cell type and tissue-enriched biomarkers, as well as candidate genes implicated in biological processes of interest for functional interrogation via whole-animal RNAi knock-down (Alvarado and Newmark 1999; Newmark et al. 2003; Reddien et al. 2005). Efficient integration and synthesis of this massive and expanding trove of visual, molecular, and functional data requires the adoption of universal standards, including a common anatomical vocabulary and syntax, and most importantly, a method of organization that allows data to be easily retrieved by any category.

Big data must be readable, reusable, and extensible by both humans and computers. Ontologies (Gruber and Others 1993) excel at this crosstalk, creating common understanding within a domain of knowledge by placing entities described in a controlled language in relationship to each other using either explicitly defined (asserted) or inferred statements. The resulting structure is a representation of knowledge, readable by both humans and machines, that is able to retrieve both asserted and inferred axioms, i.e., a knowledge graph. This structure has made ontologies ubiquitous in the digital age, where frameworks such as the semantic web facilitate information sharing across automated systems. Codifying knowledge using an ontological framework also promotes sharing data according to FAIR (Findable, Accessible, Interoperable, and Reproducible) practices (Wilkinson *et al*., 2016).

Ontology structures can be used to explore, understand and discover relationships among data and to develop testable hypotheses. A clear example of the usefulness of ontologies in the biological sciences is the Gene Ontology (GO). GO is a well-known,highly used framework that endeavors to ascribe putative functions to genes across species based on sequence homology, from the molecular to the organismal level (Ashburner *et al*., 2000). The hierarchical organization in GO defined by relationships between terms facilitates refinement or expansion of candidate gene lists from gene expression studies. For example, a gene list associated with pigmentation can be refined by selecting a more granular category such as cellular pigmentation. GO also provides a first-pass analysis of molecular and cellular processes most likely to be enriched or perturbed between experimental samples. Additionally, data can be interrogated to inform if genes are assigned to multiple biological processes or other categories.

Anatomy ontologies are another example of the use of ontologies in biology and have been developed for many well-established and emerging research organisms, including slime molds (Gaudet *et al*., 2008), nematodes (Lee and Sternberg, 2003), fruit flies (Costa *et al*., 2013), frogs (Segerdell *et al*., 2008), zebrafish (Van Slyke *et al*., 2014), mice (Hayamizu *et al*., 2013), and humans (Bard, 2012). In addition to providing a controlled vocabulary and means of streamlining data annotation, these frameworks also facilitate comparative studies on animal development and evolution. One way this is accomplished is by making species-specific ontologies compatible and interoperable with Uberon, a cross-species gross anatomy ontology (Mungall *et al*., 2012). The interoperability of ontologies enriches and extends navigation among disparate datasets. For example, it will soon be possible to identify evolutionarily conserved genes required for ciliogenesis, along with genes expressed in cilia, via searches that use GO, Uberon, and species-specific anatomy and phenotype ontologies. To maximize the scientific utility and visibility of big data generated by the planarian research community, the field requires new bioinformatic tools, including ontologies, that will improve data archival practices and search functions across different experimental platforms. Here we debut the Planarian Anatomy Ontology (PLANA) and demonstrate its utility for data annotation and integration across 155 published data sets.

## Results

### Annotating anatomical terms: Classes

PLANA is an extendable framework of defined terms that aims to holistically describe *Smed* anatomy across all life cycle stages for the asexually and sexually reproducing biotypes. In order to ensure that PLANA encompasses all relevant anatomical *Smed* terms, we conducted a review of 200 primary research citations from 2005 through 2019 (Table S1) and identified 658 terms pertaining to biotypes, life cycle stages, embryonic, adult and regenerating anatomical structures, subcellular components, cells, tissues, organs, anatomical systems, body regions, anatomical spaces (e.g., cavities and lumens), anatomical surfaces, boundaries, planes, and axes. 380 of the 658 terms were synonymous (e.g., eye and photoreceptor) resulting in a final set of 278 terms commonly used by the planarian community. Hereafter, we call these terms classes. Each class has a primary name (label) and may have supporting synonym(s). In addition, classes were imported from other ontologies and composite classes (described below) were created. In all, PLANA version v2020-07-31 has a final class count of 855.

While a list of anatomical terms is useful, the strength of an ontology derives from the ability to annotate classes with metadata and to hierarchically organize classes into a relational network. Each class has its own set of categorical, spatial, temporal and developmental relationships to other classes (Figure 1). Following the convention set forth by (Van Slyke *et al*., 2014), classes are represented using single quotation marks, and while their ID generally follows (e.g.,’epidermis’ PLANA:0000034), we omit the ID for readability. All IDs for PLANA classes mentioned in the text are found in Table S2.

**Figure 1:**
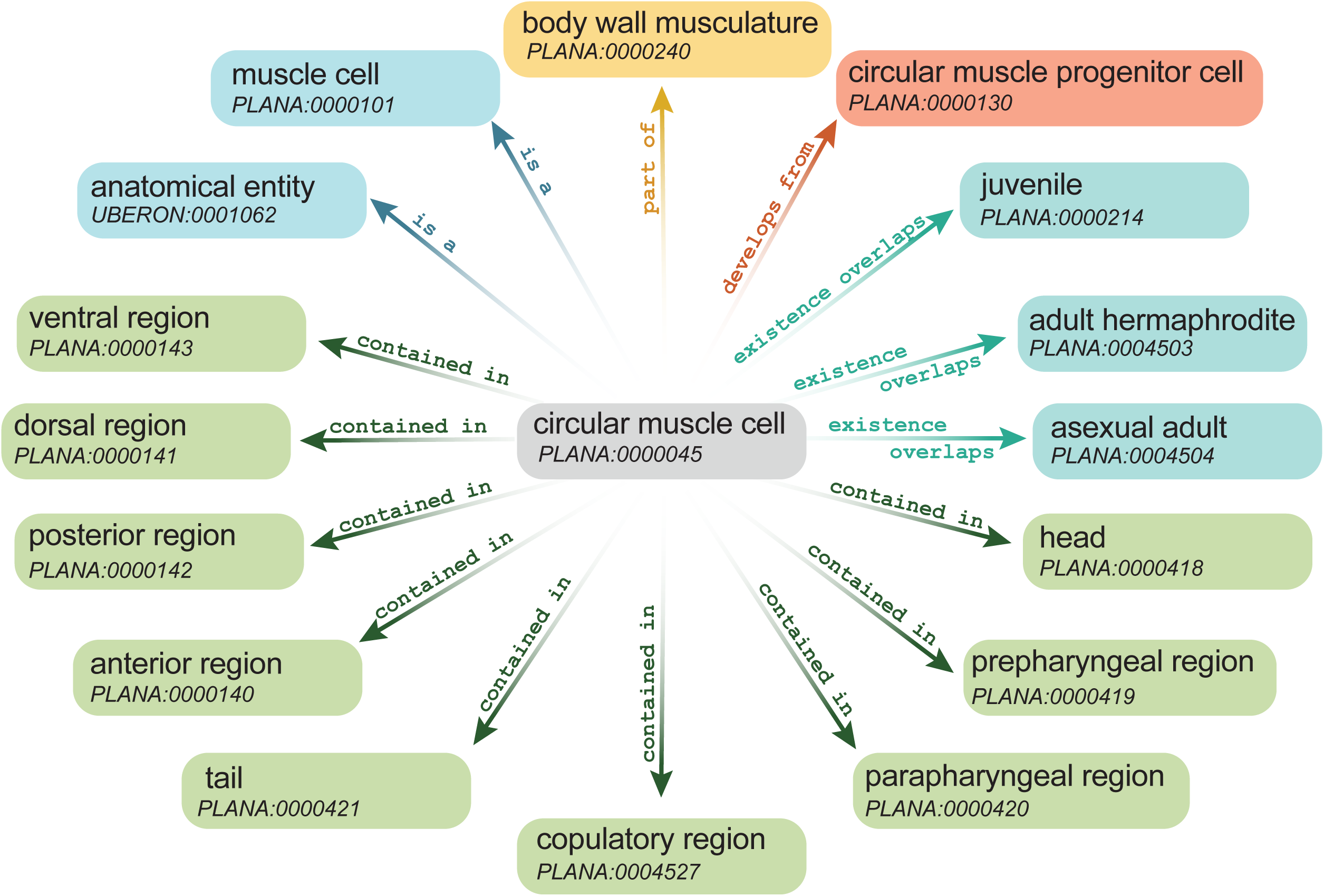
PLANA classes are linked to one another through relationship terms. The anatomical class ‘Circular muscle cell’ is shown in relation to other PLANA classes. Different colors reflect the different types of relationships between the classes. Relationships shown are: is a (blue), part of (yellow), develops from (lavender), existence overlaps (aqua), and contained in (green).

Each class was annotated with required information as follows: a singular name (label), e.g., ‘eye’, a unique identification number (ID), a definition, and the relevant reference(s) for the definition (Figure 2A). Optional annotations were also assigned to classes (Table 1), including synonyms, external ontology database identification numbers (dbxref) to facilitate comparative anatomy searches, and images depicting anatomical features, along with explanatory legends and references (Figure 2A). Together, all classes and their relationships comprise a large, self-organizing webwork (Figure 2B).

**Table 1:** Class Annotations.

| **Annotation** | **Methods for Content** | **Required/Optional** |
| --- | --- | --- |
| **label** | The primary name for a new class. When more than one potential name for a class exists, both prevalence and accuracy are considered. Alternate names appear as synonyms. | required |
| **definition** | Succinct statements, supported by published literature, codifying descriptions and salient characteristics for a class. Definitions are written by domain experts (planarian biologists). | required |
| **definition database cross-reference** | PMID or OCLC identification number for the first publication(s) that introduce and define the class. This information is located within the definition annotation. | required |
| **synonym** | Additional name(s) for a class. This field encompasses broad and exact synonyms, as well as colloquial synonyms. | optional |
| **database cross-reference** | Database identification number(s) for classes imported from other ontologies, as well as identification numbers for publications from the literature search (Table S1) that reference the class. | optional |
| **depicted by** | Image representing the class. Images published previously must be available for use under open source agreements or used with permission. | optional |
| **comment (depicted by)** | Legend that describes the image shown in the “depicted by” field. Located within the “depicted by” annotation. | optional |
| **comment** | Clarifying statement for the class outside of the definition. | optional |
| **created by** | ORCID of author. | optional |

**Figure 2:**
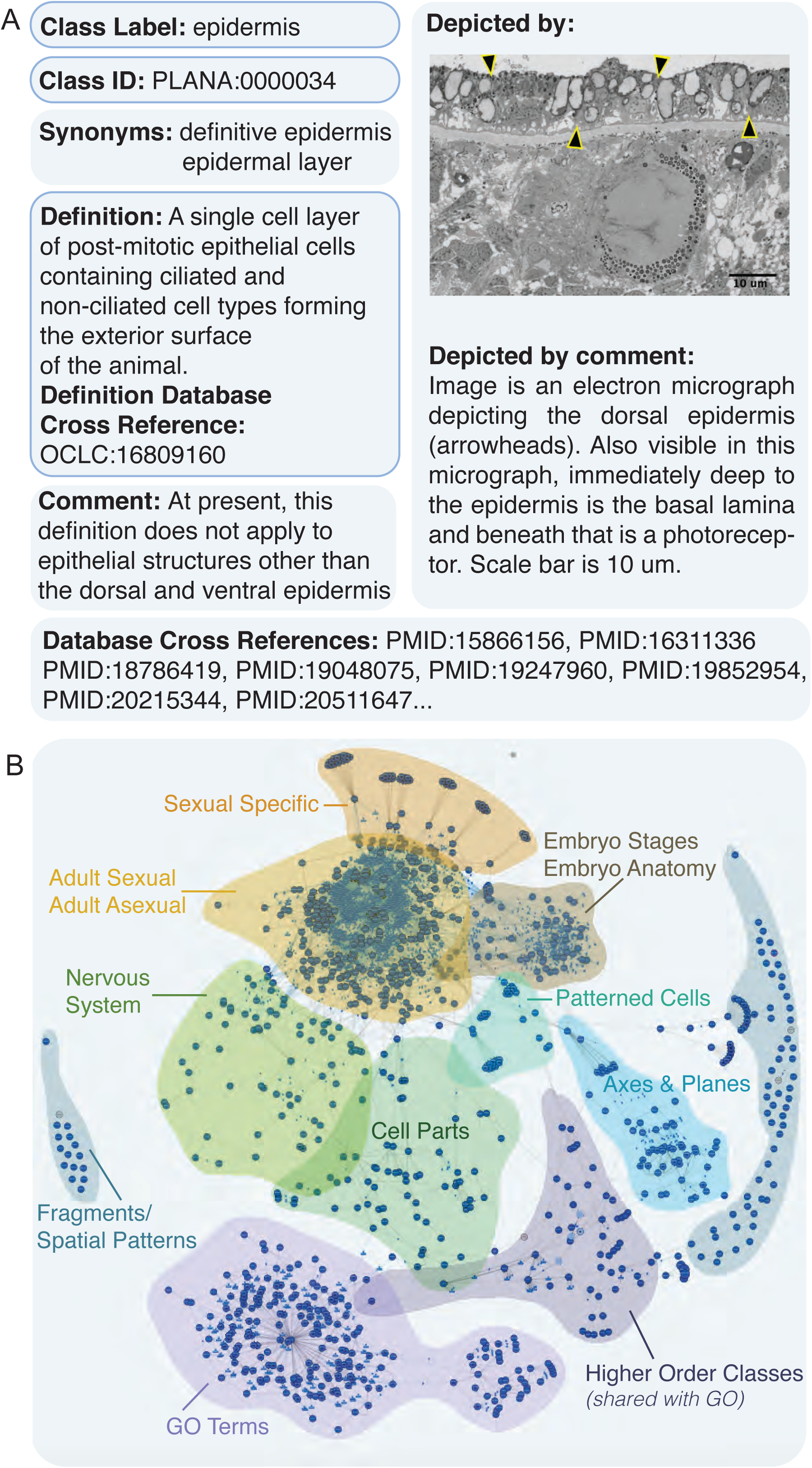
PLANA class annotation fields and structure. A) An example of required (blue outlines) and optional annotations for the class ‘epidermis’. B) WebVowl visualization of PLANA structure. Each class is represented by a dark blue point. The proximity between classes is a metric of similarity and relationships between classes (object property-based axioms). Clusters of classes are noted with their categories.

In order to extend the use of PLANA and promote interoperability with other ontologies, classes at the top of the Uberon anatomy ontology hierarchy (Mungall et al. 2012) were imported directly into PLANA (e.g., ‘anatomical entity’ UBERON:0001062 and ‘life cycle stage’ UBERON:0000105). For these imported classes, no annotation fields (e.g., ID) were altered and no additional annotations were added. Imported classes are subject to change when Uberon is updated. These imported classes frequently appear as nodes in the PLANA hierarchy (Figure 2B).

To facilitate comparative anatomy queries across species, additional classes from extant anatomy ontologies were imported and instantiated into PLANA whenever possible, including Uberon (Mungall *et al*., 2012), the Common Anatomy Reference Ontology (CARO) (Haendel *et al*., 2008), the Biological Spatial Ontology (BSPO) (Dahdul *et al*., 2014), the Cell Ontology (CL) (Diehl *et al*., 2016), and the Gene Ontology (GO) (Ashburner *et al*., 2000; The Gene Ontology Consortium, 2019). For these instantiated classes, new PLANA IDs were assigned, and annotations were edited or added to reflect planarian-specific information. For example, the class ‘eye’ UBERON:0000970 was given the ID PLANA:0000036 (the external ontology identification number was retained in the database cross-reference (dbxref) annotation field). ‘Eye’ PLANA:0000036 was annotated with planarian-specific information pertaining to its cellular origin, development, anatomical system, and anatomical location. Additional annotations that altered the original imported ‘eye’ class after instantiation included the incorporation of a representative image, as well as PubMed IDs for the use of ‘eye’ in planarian literature. Such class instantiation prevents unsanctioned changes within PLANA when external ontologies are updated yet allows association with outside ontologies via the dbxref field.

### Synonym annotation

Ontology interoperability and applications involving human input require PLANA to allow for variability in language. This variability is accommodated by annotating classes with synonyms. Synonyms make PLANA more flexible for users, enabling queries and searches to be more inclusive. When synonymous names were present in the publication record, class labels were assigned to the most commonly used term, and less frequently used names were annotated as synonyms. For example, ‘cephalic ganglia’ has the synonyms brain, cerebral ganglia, and bi-lobed brain. Classes may be annotated with multiple synonyms (Figure 2A). Exceptions to this rule include instances where classes were imported from another anatomy ontology and represented a broad comparative anatomical name. For example, ‘eye’ PLANA:0000036 (UBERON:0000970) superseded the popular moniker photoreceptor as the class name in PLANA in order to strengthen cross-species comparisons. Popular names that lacked specificity were not used as class labels, e.g., ‘epidermis’ (the outermost epithelial covering of the animal) was selected as a class name rather than the often-used term epithelium, as there are many other epithelial tissues apart from the ‘epidermis’.

### Class definitions and references annotations

In order to clarify both class structures and meaning for planarian biologists, comparative anatomists, and ontologists alike, each class has a written definition embedded with corresponding published reference(s) demonstrating the first use of the term in our literature search (def_dbxref, Figure 2A, Table S2). Original external ontology ID(s) from instantiated classes are held in a dbxref annotation field. The dbxref field also contains PubMed identification numbers for articles from the literature search that contain that class (Figure 2A, Table S1). Taken together, the class definitions and database cross-references provide provenance and promote ontology interoperability.

### Prototypic imagery

Through its definitions and links to publications, PLANA is inherently an educational resource. To increase PLANA’s didactic potential, we appended images to classes using the optional “depiction” annotation field and added explanatory legends using the “comment” annotation field (Figure 2A). Over 200 classes in this release are accompanied by an archetypal image, either an illustration for spatial classes or an electron or light microscopy image for anatomical structures. This imagery augments the written class definition through clear visualization (Figure 2A).

### Composite classes

During the literature survey to generate class names (Table S1), multi-word classes were included (e.g., ‘photoreceptor neuron’). However, the need to create more specific terms became apparent (e.g., ‘anterior photoreceptor neuron’ and ‘posterior photoreceptor neuron’). Moreover, it became clear that many classes could be created using an additive, formulaic approach already employed by other ontologies. Pre-composed, or composite classes (Mungall et al. 2010) were created using patterns that auto-generate a new class by combining two existing classes using Dead Simple OWL Design Patterns (DOSDP) (Osumi-Sutherland *et al*., 2017) (Figure 3A). Composite definitions were auto-generated and may be overwritten by curators. Furthermore, composite classes may be used to generate new classes, providing a stereotypical way to generate terms with greater specificity (Figure 3A). Patterns used to make composite classes appear in Table S3. This automated addition of classes allows rapid expansion of more specific classes as the need for spatial and temporal granularity grows.

**Figure 3:**
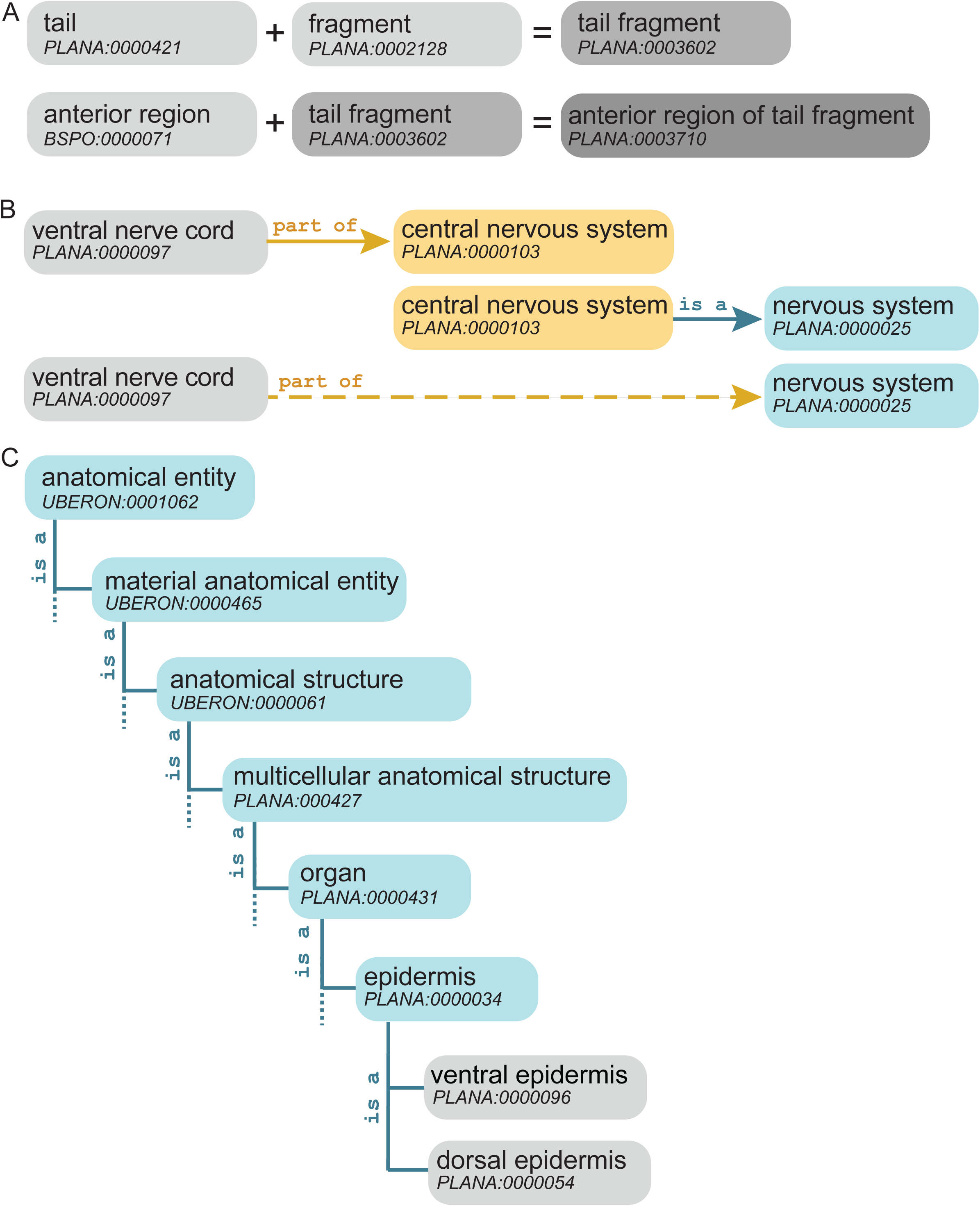
Creation of new classes using patterning algorithms and relationship. A) Composite classes, e.g., ‘tail fragment’ PLANA:0003602, generated by patterning algorithms, may be used to create new classes with greater specificity, e.g., ‘anterior region of tail fragment’ PLANA:0003710. B) Object property transitivity allows relationships to be inferred indirectly, across multiple layers of the PLANA hierarchy. Solid arrows are asserted axioms in PLANA, while the dashed arrow is an inferred relationship. C) Relationship hierarchy for the dorsal epidermis and ventral epidermis classes through the “is a” relationship.

### Constructing a relational structure: relations and object properties

An ontology’s strength lies within its hierarchical structure, which is provided by a single baseline categorical relationship (*is a*) working together with more specific relational terms called object properties. This release of PLANA uses 14 object properties from the Relationship Ontology (RO) (Smith et al. 2005) which enable the construction of categorical, spatial, developmental, and temporal relationships between classes (Table 2). Following convention, object properties are herein formatted using lowercase lettering and the font `Courier New` (Van Slyke *et al*., 2014). For example, the ‘ventral nerve cord’ is `part of` the ‘central nervous system’. Some object properties have the feature of being transitive, meaning the property can be inherited by a subclass, or entailed through the hierarchy. To expand on the previous example, `part of` is transitive; therefore, since the ‘central nervous system’ is a ‘nervous system’ and the ‘ventral nerve cord’ has been asserted as being `part of` the ‘central nervous system’, ‘ventral nerve cord’ can also be inferred to be `part of` the ‘nervous system’ (Figure 3B).

**Table 2:** Object Property Relationship Terms.

| Object Property | Realm | Transitive | Definition | Specific Rules for Use in PLANA | Example |
| --- | --- | --- | --- | --- | --- |
| **part of** | scalar<br>categorical | Yes | X *part of* Y<br>where X is a<br>more specific<br>class and Y is a<br>broader class | A core relation that holds between a part and its whole. Used in PLANA to assign cells, tissues, and organs to anatomical systems. Since these relationships are transitive, relationships are explicitly made spanning one level of anatomical organization: cell to tissue, tissue to organ, or organ to system. | 'pigment cup cell'<br>PLANA:0000031<br>part of 'optic cup'<br>PLANA:0000075 |
| **contained in** | spatial | No | X *contained in* Y where X is a more specific class and Y is a broader body region class. | Assigns the spatial location of an anatomical entity to a region of the embryo, asexual adult, or adult hermaphrodite body plan. Since the *contained in* object property is not transitive, *contained in* is annotated for all cell types, tissues, and organs within an anatomical system. | 'pigment cup cell'<br>PLANA:0000031<br>contained in<br>'anterior region'<br>PLANA:0000140 \|<br>'dorsal region'<br>PLANA:0000141 \|<br>'head'<br>PLANA:0000418 |
| **anterior to** | spatial | Yes | X <sup>anterior</sup> to Y if X is further along the antero-posterior axis than Y, towards the head | Defines relative positions of two classes along the anterior-posterior axis of the animal, where the anterior-most structure on the axis is the head tip and the posterior-most structure is the tail tip. | 'head'<br>PLANA:0000418<br>anterior to<br>'prepharyngeal region'<br>PLANA:0000419 |
| **posterior to** | spatial | Yes | X <sup>posterior</sup> to Y if X is further along the antero-posterior axis than Y, towards the tail. | Defines relative positions of two classes along the anterior-posterior axis of the animal, where the anterior-most structure on the axis is the head tip and the posterior-most structure is the tail tip. | 'prepharyngeal region'<br>PLANA:0000419<br>posterior to 'head'<br>PLANA:0000418 |
| **adjacent to** | spatial | No ;<br>Symmetric | X <sup>adjacent</sup> to Y if and only if X and Y share a boundary. | Used at cell and tissue levels when more granular spatial information other than shared boundary is known. **Not** used to describe relationships like body regions adjacent to each other (see *anterior to*, *posterior to*). | 'collecting duct'<br>PLANA:0000118<br>adjacent to 'distal tubule'<br>PLANA:0000053 |
| **immediately superficial to** | spatial | No | X <sup>immediately superficial</sup> to Y if X is further along the medio-lateral axis towards lateral than Y and X shares a boundary with Y. | Defines relative positions of two classes along the medial-lateral axis of the animal, where the lateral-most structure on the axis is the epidermis and the medial-most structure is likely the gut lumen. | 'epidermis'<br>PLANA:0000034<br>immediately superficial to 'basal lamina of the epithelium'<br>PLANA:0001005 |
| **immediately deep to** | spatial | No | X immediately deep to Y if X is further along the medio-lateral axis towards medial than Y and X shares a boundary with Y. | Defines relative positions of two classes along the medial-lateral axis of the animal, where the lateral-most structure on the axis is the epidermis and the medial-most structure is likely the gut lumen. | 'basal lamina of the epithelium'<br>PLANA:0001005 immediately deep to 'epithelium'<br>PLANA:0001005' |
| **develops from** | developmental | Yes | X develops from Y if and only if either (a) X directly develops from Y or (b) there exists some Z such that X directly develops from Z and Z develops from Y. | Describes developmental provenance of cell, tissue, organ, and organism stage classes from other cell, tissue, organ and organism stage classes. **Not** used to describe production and manufacture of biochemicals and substances (see produced by). | 'neoblast'<br>PLANA:0000429 develops from 'blastomere'<br>PLANA: 0004517 |
| **produced by** | process | No | X produced by Y if some process that occurs in X has output Y. | Describes manufacturing of any physical product that is not a cell, tissue or organ class by a cell, tissue or organ class. **Not** used to describe developmental provenance (see develops from). | 'mucus'<br>PLANA:0002059 produced by 'secretory cell'<br>PLANA:0000105 |
| **existence starts during or after** | temporal | No | X existence starts during or after Y if time point X starts ≥ time point Y starts. | Describes temporal provenance for any anatomical entity with respect to developmental stages. | 'blastomere'<br>PLANA:0004517 existence starts during or after 'Stage 1'<br>PLANA:0000001 |
| **existence overlaps** | temporal | No | X existence overlaps Y if and only if either (a) the start of X is part of Y or (b) the end of X is part of Y. | Describes range of temporal existence for any anatomical entity with respect to developmental stages. | 'blastomere'<br>PLANA:0004517<br>existence overlaps<br>'Stage 1'<br>PLANA:0000001\|<br>'Stage 2'<br>PLANA:0000002\|<br>'Stage 3'<br>PLANA:0000003 \|<br>'Stage 4'<br>PLANA:0000004 \|<br>'Stage 5'<br>PLANA:0000005 |
| **existence ends during or before** | temporal | No | X existence ends during or before Y if time point X ends <= time point at which Y ends. | Describes temporal extinction for any anatomical entity with respect to developmental stages. | 'blastomere'<br>PLANA:0004517<br>existence ends during or before<br>'Stage 5'<br>PLANA:0000005. |

### Baseline categorical relationship: is a

The categorical “is a” relationship is the baseline relationship that sets up PLANA’s class: subclass (parent: child) structure, independent of other relationships conferred through object properties. Most classes are linked to at least one other class through an “is a” relationship (e.g., ‘ventral epidermis’ is a ‘epidermis’). The “is a” relationship forms parent-child relationships between terms. Broadly defined, parent classes occupy relatively higher-order positions in the hierarchy, while child terms are more specific. In the example above, ‘epidermis’ is the parent class and ‘ventral epidermis’ is the child class (Figure 3C). A class can have multiple parents, e.g., ‘sperm’ is a ‘gamete’, and ‘sperm’ is a ‘male germ cell’.

### Scalar categorical relationship: part of

The `part of` object property codifies scalar relationships from cell type to tissue, tissue to organ, organ to anatomical system, and anatomical system to the whole organism. Taken in one step of ‘class’ `object property` ‘class’, relationships are as simple as the earlier mentioned example: ‘ventral nerve cord’ `part of` ‘central nervous system’. Like many object properties used in PLANA, `part of` relationships are transitive (Table 1; Figure 3B). As previously mentioned, transitivity enables relationships to be inferred among terms when the ontology is queried. A more complex *explicitly* asserted example of the `part of` object property has many classes chained together: ‘intestinal phagocyte’ `part of` ‘gastrodermis’, ‘gastrodermis’ `part of` ‘gut’, ‘gut’ `part of` ‘digestive system’, and finally ‘digestive system’ `part of` ‘asexual organism’. Statements that leap steps in scale, like ‘intestinal phagocyte’ (cell) `part of` ‘gut’ (organ), and ‘gastrodermis’ (tissue) `part of` ‘digestive system’ (anatomical system) are *not explicitly* asserted in PLANA but *are inferred* and are true statements. Furthermore, inferred relationships may be made among statements constructed using different object properties. For example, ‘central nervous system’ is a ‘nervous system’ and the ‘ventral nerve cord’ `part of` ‘central nervous system’, thus ‘ventral nerve cord’ `part of` ‘nervous system’ is inferred (Figure 3B).

### Spatial relationships: contained in, anterior to, posterior to, immediately deep to, immediately superficial to, adjacent to

PLANA has several object properties used to denote positional relationships among classes (Table 1). The `contained in` object property associates cell, tissue, organ, and anatomical systems with classes defining broad spatial domains of the intact embryo, juvenile, and adult body plans (Figure 4). Examples include ‘embryonic pharynx’ `contained in` ‘oral hemisphere’ (Figure 4A), and ‘photoreceptor neuron’ `contained in` ‘anterior region’, ‘dorsal region’, and ‘head’ (Figure 4C). For experimental data annotation, the `contained in` object property may also be used to associate classes with regions of regenerating adult asexual or hermaphrodite body fragments (e.g., ‘blastema’ `contained in` ‘head fragment’).

**Figure 4:**
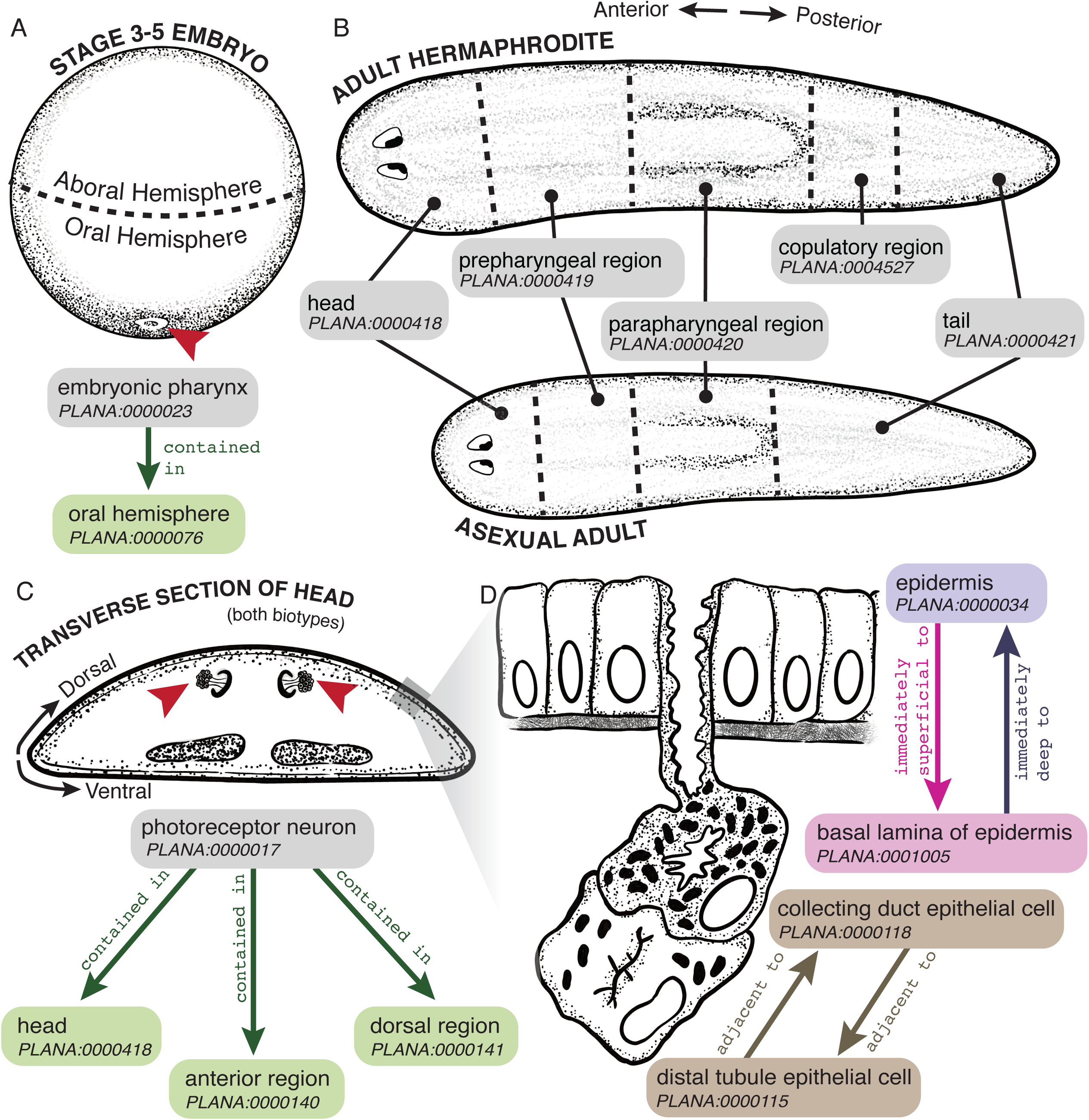
Codifying spatial relationships using the contained in object property. *Smed* embryonic (A) and adult (B) body plans. A) The ‘embryonic pharynx’ (red arrowhead) is `contained in` the ‘oral hemisphere’ of ‘Stage 3’, ‘Stage 4’ and ‘Stage 5’ *Smed* embryos. B) The body region classed for the ‘adult hermaphrodite’ and ‘asexual adult’. C) ‘photoreceptor neuron’ (red arrowheads) is `contained in` the ‘anterior region’, the ‘dorsal region’ and the ‘head’. Gray box denotes epidermal and subepidermal region depicted in D) where the ‘epidermis’ is `immediately superficial to` the ‘basal lamina of the epidermis’ which is in turn `immediately deep to` the ‘epidermis’. Another spatial relationship is that the ‘collecting duct epithelial cell’ and ‘distal tubule epithelial cell’ are `adjacent to` each other.

Spatial relationships along the anteroposterior axis are described using the `anterior to` and `posterior to` object properties. For example: ‘head’ `anterior to` ‘prepharyngeal region’. Spatial relationships along the mediolateral axis are codified using the object properties `immediately deep to` and `immediately superficial to`. For example, ‘epidermis’ `immediately superficial to` ‘basal lamina of the epithelium’, and reciprocally, ‘basal lamina of the epithelium’ `immediately deep to` ‘epidermis’ (Figure 4D). Classes next to each other in the body, but not present in a fixed position relative to the anteroposterior, dorsoventral or mediolateral body axes, may be linked using the reciprocal `adjacent to` object property. In the proto-kidneys (‘protonephridia’) the asserted axiom ‘collecting duct epithelial cell’ `adjacent to` ‘distal tubule epithelial cell’ also results in a reciprocal inferred axiom of ‘distal tubule epithelial cell’ `adjacent to` ‘collecting duct epithelial cell’ (Figure 4D)(Rink et al. 2011; Scimone et al. 2011).

### Temporal relationships: existence starts during or after, existence overlaps, and existence stops during or before

Where known, temporal staging information was annotated for classes using the `existence starts during or after`, `existence overlaps`, and `existence stops during or before` object properties. For example, ‘embryonic pharynx’ `existence starts during or after` ‘Stage 2’; ‘embryonic pharynx’ `existence overlaps` ‘Stage 2’, ‘Stage 3’, ‘Stage 4’, ‘Stage 5’, ‘Stage 6’; ‘embryonic pharynx’ `existence stops during or before` ‘Stage 6’. The `existence overlaps` object property was applied to cell types, tissues, organs, and anatomical systems present in asexual adults and/or juvenile and adult hermaphrodites.

### Developmental relationship: develops from

Developmental provenance, where known, was codified using the develops from object property. In planarians, lineages for all adult body cell types descend from a population of cycling adult pluripotent stem cells called neoblasts (Newmark and Sánchez Alvarado, 2000; Wagner, Wang and Reddien, 2011; Zeng *et al*., 2018). As neoblasts commit to a cell-type specific differentiation program, they are thought to down-regulate expression of stem cell-enriched genes, exit the cell cycle, and concomitantly upregulate expression of lineage-promoting transcription factor(s) (Guo, Peters and Newmark, 2006; Scimone *et al*., 2011; Wagner, Wang and Reddien, 2011; Shibata *et al*., 2016; Zeng *et al*., 2018). In PLANA, lineage trajectories are denoted unidirectionally, originating in the neoblast population and proceeding through one or more documented cell state transitions to a terminally differentiated cell type. A well-studied example, the epidermal lineage, is documented as follows: ‘Category 2 cell’ develops from ‘zeta neoblast’, ‘Category 3 cell’ develops from ‘Category 2 cell’, ‘Category 4 cell’ develops from ‘Category 3 cell’, and ‘Category 5 cell’ develops from ‘Category 4 cell’ (Figure 5) (Eisenhoffer, Kang and Sánchez Alvarado, 2008; Pearson and Sánchez Alvarado, 2010; van Wolfswinkel, Wagner and Reddien, 2014; Tu *et al*., 2015; Cheng *et al*., 2018). ‘zeta neoblast’ and Category 2, 3, and 4 cells are all an ‘epidermal progenitor cell’, while Category 5 cells are a ‘terminally differentiated cell’.

**Figure 5:**
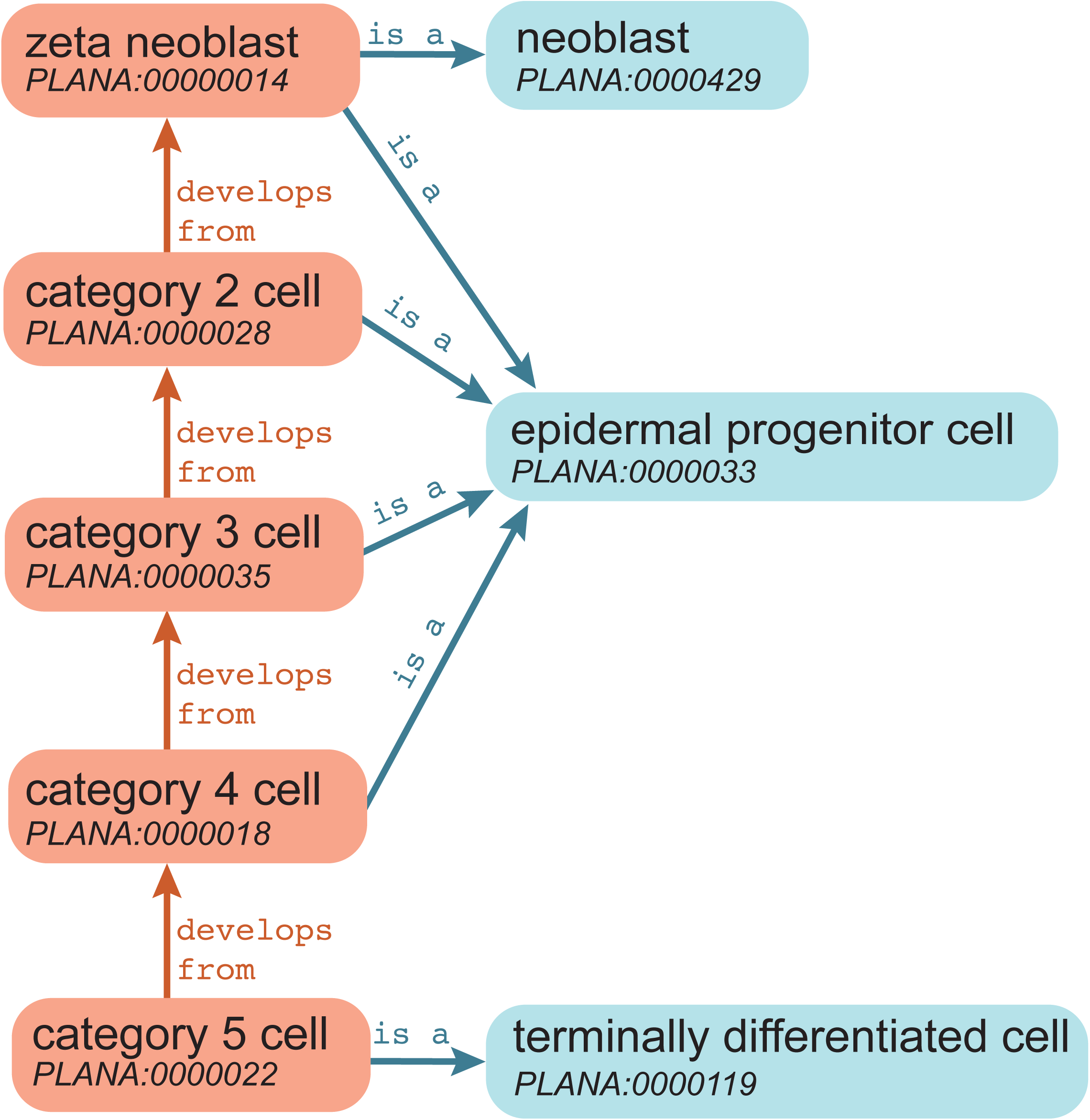
Ontogeny is recorded using the develops from object property. Schematic showing both the “is a” relationship and `develops from` object property charting a proposed lineage trajectory for the epidermal lineage, from stem cell to terminally differentiated cell type.

### Checking the structure: Queries

To ensure the veracity of asserted and inferred axioms, we systematically queried PLANA using the competency questions listed in Table S4. When a query return contained errors such as incorrect relationships between classes (e.g when ‘ovary nerve plexus’ `part of` ‘asexual adult’ was returned), asserted axioms were edited and/or added to correct the inferred error. This process was iterated until all returned classes were correct.

### Talking to other ontologies: interoperability

Bespoke, field-specific ontological frameworks are useful for data organization but become extensible and more powerful when designed to work with other ontologies. Optimal interoperability between PLANA and other ontologies was achieved by importing higher-order parent classes from Uberon and instantiating classes from other ontologies, along with recording their original ID as a dbxref annotation.

Additional interoperability is built into composite classes, as classes from the BSPO (Dahdul *et al*., 2014), GO (Ashburner *et al*., 2000; The Gene Ontology Consortium, 2019), and the Phenotype and Trait Ontology (PATO) (http://www.obofoundry.org/ontology/pato.html) were imported into PLANA upon creation of composite classes. For example, GO terms for mitotic and meiotic cell cycle phases were imported to generate PLANA composite classes for stages of the neoblast cell cycle (e.g., S phase neoblast), the mitotic germ cell cycles (e.g., metaphase spermatogonia), and meiotic germ cell cycles (e.g., meiotic metaphase 1 stage spermatocyte) (Table S3).

While direct import and instantiated use of classes from other ontologies is important for interoperability, another equally fundamental means of ensuring that one ontology can talk to another is through limiting object properties to those referenced in the Relation Ontology (RO) (Smith et al. 2005). The RO is a reference set of relations and their semantics used for standardization across ontologies in the OBO Foundry (Smith et al. 2007). Our strict use of RO object properties ensures that PLANA relationships are found in, and stated similarly, as in other ontologies. PLANA was constructed with an eye towards ontology interoperability, facilitating its application to evo-devo and comparative anatomy studies. Interoperability will also promote future extension and application of PLANA as a base framework for multiple types of data organization and will allow other ontology builds to use PLANA efficiently.

### Access PLANA

Download the latest version of PLANA through the Open Biological and Biomedical Ontology (OBO) Foundry (http://www.obofoundry.org/ontology/plana.html) or the GitHub repository (https://github.com/obophenotype/planaria-ontology).

### View PLANA

Browse PLANA on Planosphere: (planosphere.stowers.org/anatomyontology). Search the PLANA class glossary and link to class webpages (https://planosphere.stowers.org/ontology). Each class webpage contains the PLANA ID, definition and citation(s), and tools for visualizing annotated object property relationships and tables with planarian transcripts known to be expressed in each class (see below). The European Bioinformatics Institute (EMBL-EBI) Ontology Lookup Service (OLS) tree (https://www.ebi.ac.uk/ols/ontologies/plana) depicts hierarchical relationships among PLANA classes. An interactive feature, Ontology Graph, dynamically depicts user-selected relationship(s) for the class of interest in either cluster or hierarchical format and generates graphic files for download (Perez-Riverol *et al*., 2017).

WebVowl,(visualdataweb.de/webvowl/#iri=http://purl.obolibrary.org/obo/plana.owl), an interactive ontology visualization tool, may also be used for exploration and graphical depictions of PLANA.

### PLANA in Action: organization of gene expression data

As publications generate large amounts of data, there is an increasing need to make this data available and searchable in centralized locations. Planosphere is an online resource aggregator for published *Smed* datasets. We demonstrated PLANA’s utility for organizing and mining large datasets by applying PLANA to the organization of an embryonic staging series and a molecular fate mapping atlas on Planosphere. Each PLANA class has its own web page on Planosphere, ensuring seamless integration of the PLANA hierarchy and class metadata into these resources (Figure 6A).

**Figure 6:**
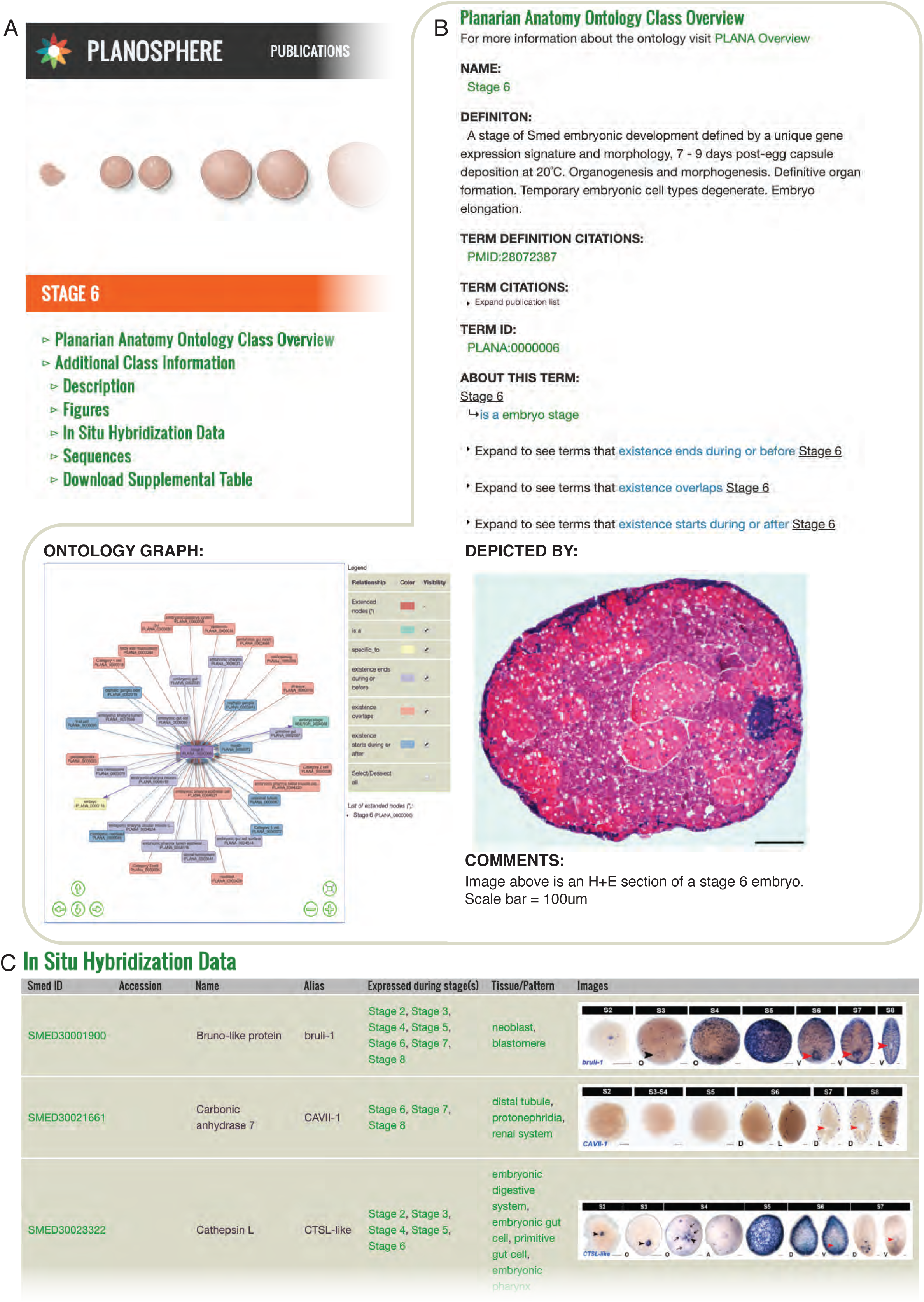
PLANA was used to create web-based resources for *Smed* embryogenesis. A) Overview of didactic tools for *Smed* embryogenesis that rely upon PLANA for organization and presentation of the data. B) Staging Resource overview. Webpage for ‘Stage 6’ PLANA:0000006 displays PLANA metadata and the Ontology Graph visualization tool. (C) Whole-mount *in situ* hybridization data was annotated and organized using PLANA classes.

### Educational resources for planarian embryogenesis

Planosphere hosts two tools powered by PLANA for exploring *Smed* embryogenesis: a staging series (https://planosphere.stowers.org/staging) and a molecular fate mapping atlas (https://planosphere.stowers.org/atlas) (Davies *et al*., 2017). The staging series defines and describes the eight stages of *Smed* embryogenesis, incorporating single embryo RNA-Seq gene expression data along with chronological and morphological information. The molecular fate mapping atlas documents cell and tissue types unique to early embryonic stages, as well as the development of adult anatomical systems. Published gene expression data from the single embryo RNA-Seq developmental time course and whole-mount *in situ* hybridization experiments on staged, wildtype embryos were annotated using PLANA. For the staging series, transcripts with enriched expression at each stage were annotated with relevant PLANA class(es) (Stage 2 - Stage 8). For the fate mapping atlas, PLANA classes for the biotype, life cycle stage(s), and anatomical structure(s) positive for expression were linked to transcripts (Figure 6B). Use of PLANA to curate gene expression data enables users to search by primary sequence, transcript identifier/name, developmental stage, and anatomical site(s) of expression, from cell type to anatomical system. Hyperlinks facilitate rapid navigation to transcript webpages (Transcript Pages) and PLANA class webpages (Figure 6C), enabling users to hone or broaden their queries, and to access relevant background information concerning embryo anatomy and development.

### Planarian Anatomy Gene Expression (PAGE)

We used PLANA to create the Planarian Anatomy Gene Expression (PAGE) database, a web-based resource that allows users to mine published gene expression data using ontological inference and PLANA classes (https://planosphere.stowers.org/search/page/about; Figure 7A). Our PAGE web forms enable users to do complex searches by term, transcript, or publication that would traditionally involve extensive literature research and elaborate manual documentation. Tasks such as identifying all transcripts expressed, across transcriptomes and research laboratories, in any structure that is a `part of` the ‘central nervous system’, or all the structures a single transcript or a group of transcripts have been published as being expressed in, now takes seconds.

**Figure 7:**
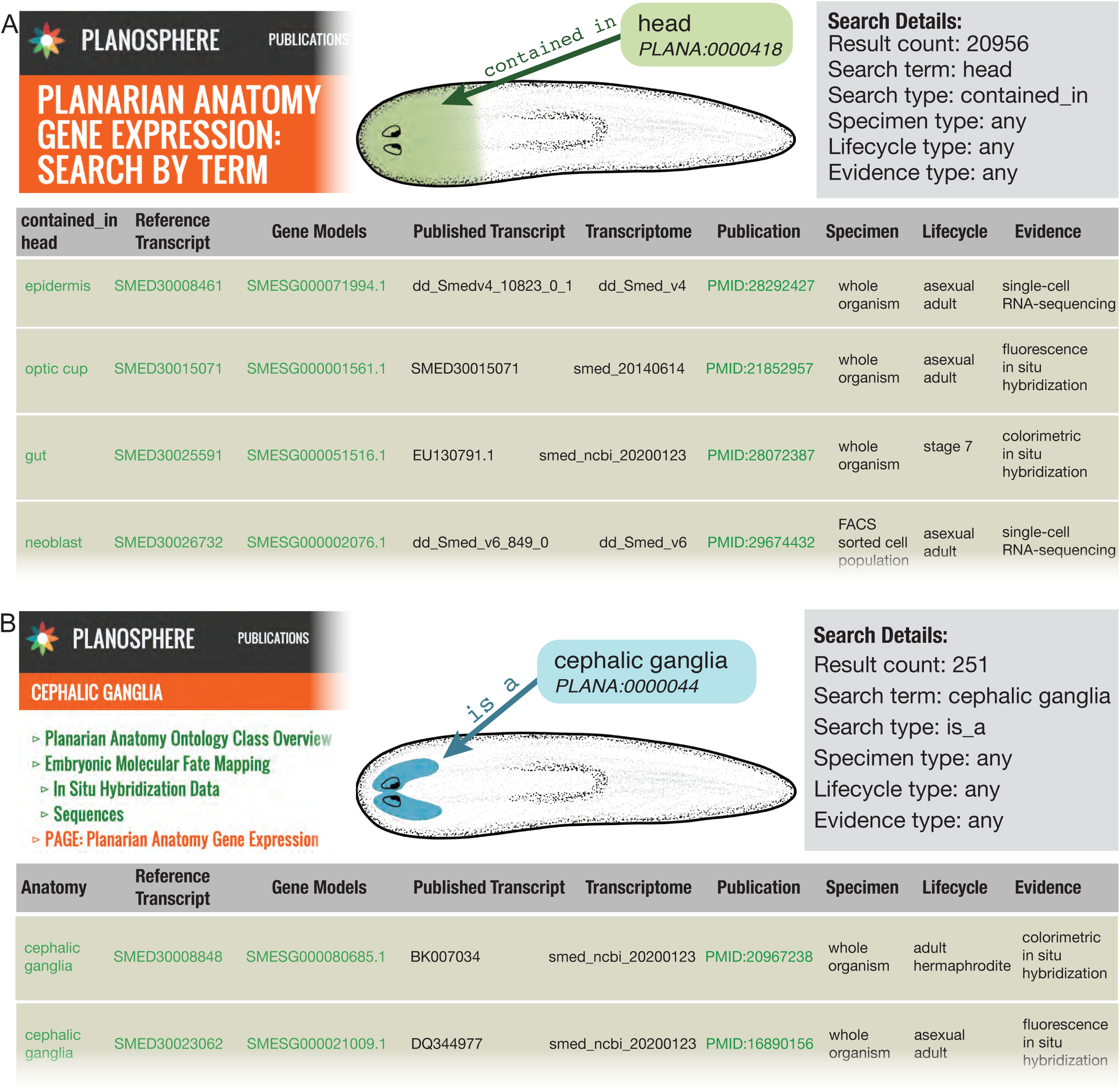
Planarian Anatomy Gene Expression Database. (A) The PAGE resource is accessible via the Planosphere website and returns a downloadable table for searches such as: find all transcripts annotated as expressed in anatomical structures contained in the head across all lifecycle stages, specimen type and evidence types. Search by transcript or publication not shown. (B) PAGE is incorporated into individual class webpages under the PAGE: Planarian Anatomy Gene Expression section. For example, the Cephalic Ganglia web page includes references, genes, and transcripts that are annotated as being expressed in an anatomical structure that is a cephalic ganglia.

To generate the PAGE database we curated qualitative expression data from 155 publications cited in the literature review (Table S1) to collect the following information: gene name(s), transcript identification number(s), Genbank accession number(s), PubMed identification number for the citation, evidence classes from the Evidence and Conclusion Ontology (ECO) (Chibucos *et al*., 2017; Giglio *et al*., 2019) (e.g., ‘colorimetric *in situ* hybridization evidence’, ‘fluorescence *in situ* hybridization evidence’, ‘RNA-sequencing evidence’, ‘single-cell RNA-sequencing evidence’ and ‘cDNA to DNA expression microarray’), PLANA class(es) describing anatomical site(s) of expression, and curator ORCID. In total, 88,870 instances of expression from wildtype, intact animals and sorted cell populations were manually curated in the PAGE database. Expression data in regenerating body fragments and in RNAi knock-down animals were not collected.

Because accessions and identifiers for annotations came from multiple transcriptomes and other sources like ESTs, we built a translation tool, Rosetta Stone Transcript Mapper, to map all sequences back to the smed_20140614 reference transcriptome (https://planosphere.stowers.org/search/rosettastone/blaze; Suppl Figure 1)(Tu *et al*., 2015). Using Rosetta Stone Transcript Mapper, the 88,870 annotations referenced 30,715 unique accessions. Those accessions mapped to 16,657 transcripts in the reference transcriptome, which are associated with 15,513 gene models (Grohme et al. 2018). PAGE is searchable by any anatomical term in PLANA (including synonyms), by transcript or accession number, and by publication. Using the PAGE resource, a researcher with a broad interest in transcripts annotated with a PLANA class that is contained in the ‘head’ would retrieve a downloadable list of 20,956 instances of expression data from 123 different publications; spanning 7 life cycle stages; 5 types of evidence; 44 PLANA classes; 15 different published transcriptomes; and 8,944 unique reference transcripts, which are associated with 7,473 gene models (Figure 7B). Alternatively, a researcher with a narrow interest in a specific transcript looking for all papers with documentation of expression can search PAGE by transcript, such as “dd_Smed_v6_76069_0_1”. This search returns a set of 6 homologs from 4 different transcriptomes and 7 publications. All of the homologs are described as *ovo* and documented by 3 evidence types as being expressed in 6 anatomical structures in a sliding scale of specificity from ‘photoreceptor neuron’ to ‘eye cell’ to ‘head’ (Table S6). All of these classes are `part of` ‘eye’ and thus `contained in` ‘head’.

### PLANA in the Future: New Contributions and Versions

PLANA is a living resource curated by the manuscript authors. New releases are automatically scheduled for weekly pick-up by Open Biological and Biomedical Ontology (OBO) Foundry (Table S5). PLANA will be versioned following substantive changes to the structure or monthly to pick up small changes. Queries (Table S3) will be performed for quality control prior to the release of each new version.

Members of the research community are encouraged to assist with PLANA curation through submission of a new class(es) and/or proposing edits to an existing class(es) using the GitHub issue tracker (https://github.com/obophenotype/planaria-ontology/issues). New class submissions require a class name, definition, PMID or DOI numbers for publication(s) referencing the definition, and a contact name and email address for the contributor. Two curators will review new classes and other proposed edits and will correspond with the contributor to resolve outstanding questions prior to updating PLANA. Bulk requests for new classes should be submitted using the spreadsheet template posted on the PLANA GitHub issue tracker.

The PLANA GitHub repository issues page contains a searchable history of questions and resolutions to issues raised by curators and community members. Questions may be submitted via opening a new issue to ensure the discussion and decision-making process is open, transparent, and archived. Requests to deprecate class(es) should be made by opening an issue. Obsolete classes remain visible in future versions of PLANA as deprecated classes. When a class is superseded by a new class, the deprecated class is listed as a synonym for the new class.

### Reporting

PLANA is described according to the Minimum Information for the Reporting of an Ontology Guidelines (Matentzoglu *et al*., 2018) (Table S7). PLANA is supported by the Sánchez Alvarado Lab at the Stowers Institute for Medical Research in Kansas City, Missouri, and data are licensed under a Creative Commons BY-NC 2.0 License. When using PLANA, please report the date(s) and/or version number(s) for the relevant PLANA files.

## Discussion

The planarian research community is generating transcriptomic, genomic, and phenotypic data at an unprecedented rate that is already well past the limiting amount of raw material human brains can hold, let alone infer information from. While we use databases to tackle the problem of the information quantity, these databases cannot infer attributes based upon known relationships. To mimic what the human brain does so well (quickly infer relationships among categories that are made by binning according to properties), we created an ontology framework to organize and facilitate inferential searching of anatomy related data. We created the PLANA ontology to address three critical needs in our field: 1) a primer for researchers to become familiar with an emerging research organism, 2) a controlled anatomical vocabulary, and 3) standardization of data curation, thus promoting searchability within and among large data sets. As a set of living data, PLANA also provides a platform for growth and adaptation within the field. Additionally, the design and workflow used to construct PLANA provides a guide for creating anatomy ontologies for those who find themselves facing the same problem of exponential data growth.

Searchable through the OLS and Planosphere, PLANA is an educational resource that enables users to familiarize themselves with *Smed* anatomy, life cycle stages, prototypical images, and relevant publications. We provide an example of PLANA’s utility and versatility as a data organization tool by using it to organize a staging series and fate mapping atlas for *Smed* embryogenesis. In addition, we created the PAGE community expression resource, a database that associates PLANA classes with an integrated reference for *Smed* transcripts and gene models that readily allows users to assess equivalency and make connections for spatial expression patterns and digital gene expression data produced across different platforms. We anticipate PLANA and the PAGE database will be used to assign cell or tissue identities to single-cell RNA-Seq cluster data. Using PAGE, it will be possible to quickly ascertain whether whole-mount *in situ* hybridization data has been reported for cluster-enriched biomarkers. By using PLANA and PAGE in conjunction with Seurat or UMAP-generated projections, predictive statements regarding anatomical identity may be made based on the proximity of cell clusters in expression space. In the near-term, PLANA is also being used to annotate high-resolution anatomical data from serial blockface scanning electron microscopy datasets.

As a standard for anatomical information, PLANA does not claim to be comprehensive or exact. On the contrary, we expect and welcome additions and curation from the greater scientific research community. Importantly, PLANA does not include processual, functional, or phenotypic information. PLANA does not encompass anatomy from planarian species other than *Smed*, but may be cloned, instantiated and edited to rapidly generate an anatomical ontology for other planarian species. PLANA will be instrumental to the construction and development of additional community resources, such as a planarian phenotype ontology. Notably, PLANA will facilitate the incorporation of a phenotype ontology into Upheno and Monarch (Shefchek *et al*., 2020), a semantic-based integrative data platform that connects expression and phenotypes with genotypes across species. Interoperability of PLANA with other anatomy ontologies, through Uberon, and of a *Smed* phenotype ontology with other phenotype ontologies, through Monarch, will facilitate comparative anatomy queries and cross-species genotypic and phenotypic comparisons.

## Materials and Methods

### PLANA Construction

PLANA content was amassed through the review of 200 publications (Table S1) to ensure comprehensive coverage of all anatomical entities reported by the planarian research community. Terms determined to be synonyms were annotated as “exact synonyms” rather than “broad synonyms” for clarity. All data were entered into shared Google spreadsheets. WebProtégé, because of its ease of use, and Google Docs because of its collaborative properties, were used with an initial draft version of the ontology to flesh out the underlying structure (Tudorache *et al*., 2013). Where possible, extant classes were imported from other ontologies and instantiated in PLANA.

All tools used or generated for this manuscript that have a repository or a website are cataloged in Table S5. PLANA was initialized and is maintained with the use of the Ontology Development Kit (ODK; Table S5). ODK sets up the directory and file structure and provides scripts to manage and maintain an ontology. It integrates Dead Simple OWL Design Patterns (DOSDP)(Osumi-Sutherland *et al*., 2017) for generating terms using patterns and ROBOT (Jackson et al. 2019) (Table S5) for handling imports from other ontologies, file format conversions, and validations. DOSDP uses yaml formatted patterns (Table S3) to generate similarly structured classes like, ‘testis cell’, ‘eye cell’, ‘pharynx neuron’, and ‘pharynx muscle cell’. These patterned terms were generated by combining two existing classes: an anatomical structure, e.g., ‘testis’, ‘eye’, ‘pharynx’, and a cell type e.g., ‘cell’, ‘neuron’, and ‘muscle cell’. Patterns may also specify that a class needs a name, definition, reference, and synonym. PLANA uses yaml patterns to manage all PLANA classes, dynamically pulling data from Google spreadsheets.

Protégé was used for visual inspection of the ontology and to query the PLANA structure (Musen and Protégé Team 2015). Queries were run using Protégé’s DL Query with the ELK 0.5.0 reasoner (Table S5) to ensure all terms are logically related and that no errant relationships were inferred after construction of our asserted hierarchy.

### Rosetta Stone Transcript Mapper

The publications entered into the PAGE database (Table S1) used several different transcriptome and gene identifiers. In order to unify this dataset, it was necessary to map the various identifiers to each other. To create this map we selected 10 transcriptomes available through Planmine (Rozanski *et al*., 2019), *Schmidtea mediterranea* nucleotide sequences from the NCBI (NCBI Resource Coordinators, 2016) and dd_Smed_v4 (an older version of the dd_Smed_v6 transcriptome available on Planmine)(Table S8). Sequences from all transcriptomes were aligned with blat (-minScore=100 -minIdentity=95) (Kent, 2002) to the most recent gene model transcripts (dd_Smes_v2) and to smed_20140614 (Tu *et al*., 2015). Aligned sequences were assigned to the corresponding gene models. Microarray probe sequences were aligned to reference sequences with blat (-minScore=30 -minIdentity=95) (Figure S1).

Some publications used different name formats even when using the same transcriptome. In order to address this complication, alternative transcript names were assigned (Table S9). NCBI protein accessions and names were assigned based on their corresponding NCBI nucleotide accession. Unigene identifiers from SmedGD (Robb, Ross and Sánchez Alvarado, 2008) were assigned based on their correspondence to dd_Smed_v4 identifiers, which were used in their construction. miRNAs were flagged based on sequence names and correspondence to mirBase (Griffiths-Jones, 2006; Kozomara, Birgaoanu and Griffiths-Jones, 2019). 34,864 of 35,761 unique identifiers were mapped to the reference sequence database. We have named the database of mappings and the tools to look up various IDs the Rosetta Stone Transcript Mapper.

### PAGE Construction

A web-based application for collecting planarian class annotations was built using R and Shiny (Bunn A 2013; Chang et al. 2017). R package ontologyX (Greene, Richardson and Turro, 2017) was used for traversing the ontology tree. R packages jsonlite (Ooms, 2014) and tidyverse (Wickham *et al*., 2019) were used for data manipulation (Table S5).

Publications (Table S1) were split among 3 curators to document accession numbers of transcripts and to associate expression data with PLANA anatomical structure classes. Care was taken to use anatomical terms or synonyms from the description provided in the text. Where text description was not provided or a term was not found in the ontology, the term was either added as a class or a synonym or curators picked the most relevant term present in PLANA. For example, PLANA does not contain “Cathepsin positive cell” as it is currently unclear what the exact physical anatomical structure corresponding to this state is, but as these cells are located in the parenchyma we designated all mentions of “Cathepsin positive cells” as ‘parenchymal cell’ (PLANA:30003116)(Fincher et al. 2018). For all digital expression data we relied on the decisions of the authors as to cutoff and enrichment.

Annotations were reviewed, typos identified and corrected, sequence IDs manually assigned if not computationally identifiable from the manuscript text, and all sequences mapped using the Rosetta Stone Transcript Mapper (Table S5). Sequence descriptions for the reference sequences and gene models were assigned. For smed_20140614, priority was given to Genbank descriptions (Benson et al. 2005). If Genbank descriptions were not available they were generated using AHRD (Table S5). Descriptions for dd_Smes_v2 transcripts and gene models were downloaded from Planmine using the intermine query builder (Rozanski et al. 2019).

The annotations, mappings, and sequence descriptions were organized into a triple store (Dingley 2003)(Table S5) and converted to turtle formatted files (ttl). The triple store was structured using Open Biomedical Association (OBAN) principles (Sarntivijai et al. 2016). The ttl files (annotations, mappings, descriptions), along with the PLANA ontology, Evidence and Conclusion Ontology (ECO) (Chibucos et al. 2017) owl files were loaded into a blazegraph datafile(Table S5), or journal (jnl) using blazegraph-runner(Table S5). We have Blazegraph running in a Docker (Merkel 2014) container that is web accessible to our planosphere web server. The Docker file was based on the lyrasis/blazegraph docker file (Table S5). Modifications were made to import our PAGE specific jnl and to change the name of our blazegraph instance to PAGE.

The PAGE webform searches generate SPARQL queries (SPARQL 1.1 Query Language) from the user input data. To ensure that users can only input a PLANA term, a modified version of the OLS autocomplete widget was used (Table S5). To allow SPARQL queries to incorporate the transitivity of the PLANA Ontology hierarchy and relationships using the ELK reasoner, we also run phenoscape/owlery (Table S5) through our customized docker container planosphere/owlery-plana(Table S5). Owlery is a collection of REST web services that enable querying with an OWL reasoner and a configured set of ontologies (Table S5). Through Owlery, a SPARQL query generated from our PAGE web form which asks to find all transcripts annotated as being expressed in the ‘nervous system’ (asserted) is expanded to include its transitive relation classes like ‘central nervous system’ and ‘peripheral nervous system’ (inferred) and also generates a new SPARQL query. This second SPARQL query is then used to query the jnl housed in our Blazegraph server.

### Animals and Imagery

*Smed* anatomical descriptions were based on CIW-4 asexual and sexual adults (Newmark and Sánchez Alvarado 2002). Illustrations were made using Procreate (https://procreate.art/) and Adobe Illustrator (https://www.adobe.com/products/illustrator.html). Hematoxylin and eosin (H+E) stained histological sections were prepared for CIW-4 asexual adults (Adler *et al*., 2014) and embryos (Davies *et al*., 2017), and images were acquired on a Olympus America Slide Scanner. Many prototypical images were produced from TEM, STEM, and SBF-SEM datasets of C4 asexual animals. Images were acquired on a Zeiss Merlin SEM with a STEM detector and Gatan 3View 2XP, or a Thermo Fisher Scientific/FEI Tecnai G2 Spirit BioTWIN with Gatan UltraScan 1000 CCD camera. For TEM and STEM imaging animals were prepared as in Cheng *et al.* (2018). For SBF-SEM animals were fixed as for STEM samples with *en bloc* staining steps per (Tapia *et al*., 2012; Hua, Laserstein and Helmstaedter, 2015) as follows: reduced osmium incubation was performed overnight at 4 C, TCH incubation at 40 C for 45 minutes, incubation in 1% UA overnight at 4 C and transferred to 50 C for 2 hours, and lead acetate incubation for 2 hours at 50 C. Animals then were dehydrated and infiltrated as for the STEM samples using either a hard formulation of Spurr’s resin (EMS) or Hard Plus resin (EMS). Fiji (Schindelin *et al*., 2012) was used for final adjustments and bilinear downsizing to the max dimension of 512.

### Accession Numbers

Accession numbers of transcriptomes, microarrays and additional resources used to construct the Rosetta Stone Transcript Mapper and PAGE resource can be found in Table S5. In addition, we downloaded every *Schmidtea mediterranea* sequence in the NCBI Genbank (Benson et al. 2005) nucleotide database on January 23, 2020.

## Acknowledgments

We thank Matthew Horridge and the whole Protégé course at the Stanford Center for Bioinformatics Research, Jim Balhoff for assistance with Owlery and Blazegraph, Andrew Koebbe for help with Blazegraph Docker, Mary Penne Mays for help with webform CSS and Dustin Dietz for system administration tasks. We would also like to thank Alice Accorsi, Blair Benham-Pyle, Biff Mann, and Aubrey Kent for insightful comments on the manuscript. Additional thanks to the SIMR Histology Core for slide sections and H+E staining, and to the SIMR Microscopy Core for scope training and slide scanner workflows.

## Data Availability

|  |  |
| --- | --- |
| PLANA<br>*At the time of publication: PLANA Release 2020-07-31*<br>(DOI:10.5281/zenodo.3969051) | https://github.com/obophenotype/planaria-ontology |
| PAGE curations and codebase | https://github.com/planosphere/PAGE |
| Rosetta Stone Transcript Mapper | https://github.com/planosphere/RosettaStone |
| Code for PAGE on Planosphere | https://github.com/planosphere |

All links to developed or referenced repositories are available in Supplemental Table 5. Original data underlying this manuscript can be accessed from the Stowers Original Data Repository at http://www.stowers.org/research/publications/libpb-1530.

